# Substrate binding and channeling allosterically modulate the interactions within the AlkB-AlkG electron transfer complex

**DOI:** 10.1101/2025.06.23.661152

**Authors:** Karolina Mikulska-Ruminska, Matthew Licht, Mehmed Z. Ertem, John Shanklin, Qun Liu, Ivet Bahar

## Abstract

The transmembrane alkane monooxygenase AlkB and rubredoxin AlkG form an electron transfer complex that hydroxylates terminal alkanes to produce alcohols. The recent cryoEM study of *Fontimonas thermophila* AlkB-AlkG complex (FtAlkBG) revealed its architecture, including a dodecane (D12) substrate in the diiron active site. However, the molecular mechanism of action of FtAlkBG remains unknown. Here, we examined the FtAlkBG dynamics and interactions by multiscale computations, including molecular dynamics (MD) simulations, elastic network model (ENM), and QM/MM study of the oxidative mechanism at the catalytic site of AlkB. D12 remained stably bound to the catalytic site during MD runs, coordinated by hydrophobic residues I267, L263, L264, and I133, critical to substrate stabilization. MD simulations further showed that D12 could exit the catalytic site via a well-defined translocation pathway gated by I54, S49, and F46, through two intermediate states before nearly leaving the protein, and diffuse back to the active site. Several salt bridges and conserved AlkB-AlkG contacts were enhanced upon substrate binding. Substrate binding also mediated the association/dissociation events required for efficient electron transfer. It promoted tighter coupling between the diiron center in AlkB and the iron in AlkG. The allosteric effects relevant to enzymatic activity regulated by the substrate binding and channeling were further delineated by ENM analysis which confirmed a strong coupling spanning the site of entry from the membrane, the catalytic site and the AlkB-AlkG interface. Our study provides new insights into key sites that could be targeted for developing AlkB-variants with desirable alkane conversion functions.

## Introduction

Alkanes are saturated hydrocarbons that constitute 20 to 50% of crude oils. Due to their abundance and cost-effectiveness, alkanes are attractive starting materials to produce high-value products [1]. They contain carbon and hydrogen atoms only with fully occupied electronic orbitals for their C-H bonds, making them almost chemically inert. To derivatize alkanes, they must first be activated to form alcohols, aldehydes, carboxylic acids, or epoxides, which represent a family of high-value products and feedstocks for further synthesis [2, 3].

Alkane 1-monooxygenases (AlkB) are membrane-bound non-heme diiron-containing metalloenzymes that oxidize alkanes of various lengths from C3 to C40 to produce corresponding alcohols [4–8]. AlkB was originally identified in the bacterium *Pseudomonas oleovorans,* which thrives in oil-rich environments where it can use alkanes as its sole carbon and energy source [9–12]. The OCT [13] plasmid of *P. oleovorans* contains nine genes involved in alkane metabolism, including AlkB, AlkG, and AlkT (see below), which facilitate the effective capture and conversion of alkanes to support their growth [8–10, 12, 14].

The first and key step in alkane utilization is alkane hydroxylation by a membrane-bound hydroxylase complex composed of three proteins: the membrane-bound monooxygenase (AlkB), a soluble rubredoxin (AlkG) composed of two rubredoxin domains, each containing a Fe-sulfur cluster, and a soluble rubredoxin reductase (AlkT). AlkB catalyzes the oxidation of the terminal methyl group of alkanes to produce the primary alcohols [8, 10, 12, 14–17]; AlkG transfers electrons from AlkT to the diiron center of AlkB; and AlkT is a flavin adenine dinucleotide (FAD)-dependent rubredoxin reductase that transfers electrons from NADH to the Fe-sulfur cluster of AlkG, which relays individual electrons to AlkB to enable biocatalysis [12, 18].

Recently, two cryoEM structures of AlkB from *Fontimonas thermophila* were resolved. One of them, determined by us at a resolution of 2.76 Å, contained a natural fusion of AlkG at the C-terminus of AlkB, thus forming the FtAlkBG complex [19]. The AlkB in the FtAlkBG complex had a dodecane (D12) substrate near the diiron center, and AlkG had a Fe-4S cluster. AlkG is bound to the cytoplasmic surface of AlkB, with a distance suitable for electron transfer. The second structure was determined at a lower resolution and contained AlkB only (FtAlkB) [20].

In the solved cryoEM structures, the substrate D12 occupied a long, narrow channel extending from the diiron-center active site to the middle of the transmembrane helices TM1 and TM2, suggesting that the hydrophobic substrate might enter the active site by first diffusing into the membrane bilayer. However, D12 was copurified with the FtAlkBG complex from the expression host before resolving the cryo-EM structure. Thus, the molecular mechanisms of substrate entry and dynamics are elusive. In addition, the two irons in AlkB, designated as Fe1 and Fe2, are 6.2 Å away from each other, stabilized through coordination by 5-6 conserved histidines. These irons are 13.6 and 17.1 Å away from the Fe-4S (or Fe3) cluster in AlkG, suggesting a direct electron transfer. The AlkB-AlkG interface harbors positively and negatively charged residues. It was previously hypothesized that these charged residues may be engaged in electrostatic interactions that favor electron transfer [19]. However, it is still unclear how AlkB and AlkG undergo coupled rearrangements and how these interactions affect the structure of the diiron center and the translocation of the substrate.

In this study, we investigated the coordinated motions of the FtAlkBG complex during substrate translocation using a combination of molecular simulations at multiple scales schematically summarized in **Figure 1**. We identified the key residues that mediate the entry and exit of substrate D12 into or from a hydrophobic tunnel connecting to the catalytic site (CS). The study points to the role of selected charged residues at the interface between AlkB and AlkG, and to the allosteric coupling between the binding and channeling of the substrate and the interactions at the AlkB-AlkG interface, accompanied by local structural rearrangements in the diiron-center. Our computational analysis provides insights into the intrinsic structural dynamics encoded by the topology of contacts, both intramolecular and intermolecular, within the electron transfer complex, and points to key sites and interactions whose perturbations could alter, if not impair, the cooperative motions that underlie the monooxygenase catalytic activity.

**Figure 1.**
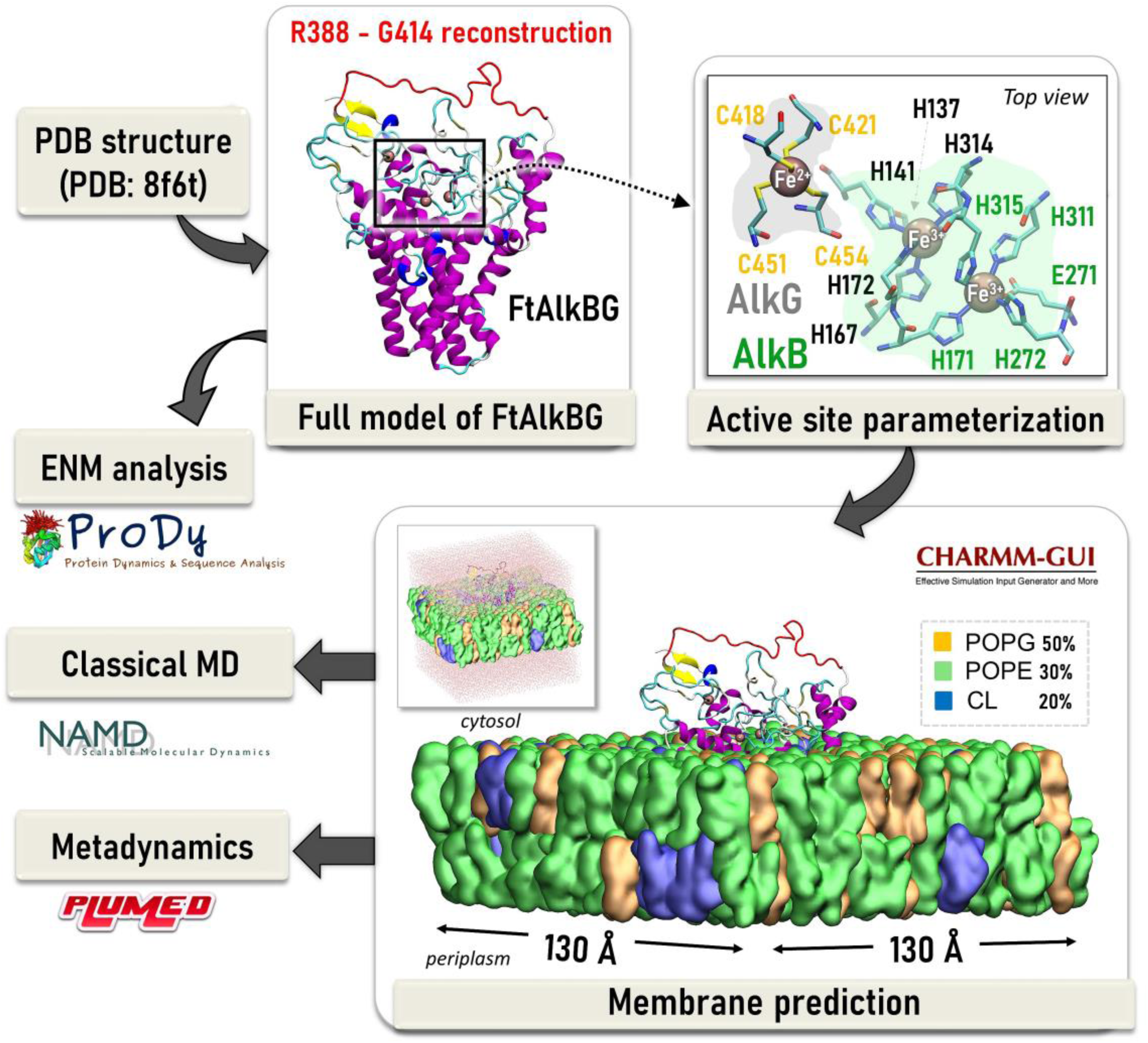
Summary of computational modeling and multiscale simulations performed for FtAlkBG. Starting from the completion of the missing components of the cryo-EM structure, parameterization of atoms at the active state using QM/MM computations, construction of the system embedded in the lipid bilayer using CHARMM-GUI [21], and two groups of simulations – classical MD and Metadynamics using the packages NAMD [22] and PLUMED [23] respectively. Additionally, elastic network model (ENM)-based analyses [24–26] were performed using the *ProDy* [27] interface. See details in the Methods.

## Results

### MD simulations point to key residues involved in stabilizing the substrate D12 at the catalytic site

To gain insights into the molecular mechanism of function of AlkB complexed with AlkG, we performed all-atom MD simulations of *Ft*AlkBG using the cryoEM resolved structure. First, the missing linker between AlkB and AlkG was reconstructed, and the enzyme was inserted into the membrane (**Fig. 1**). We used a membrane composition similar to that of *E.coli*, with 50% POPG, 30% POPE and 20% of cardiolipins (CL) [28]. After parametrization of each catalytic site to assign partial charges and geometries based on QM/MM simulations (see Methods), we performed MD simulations, in triplicate, for two systems: (i) in the presence of D12, and (ii) in the absence of D12. Of three MD runs of 200 ns performed in the presence of D12 substrate (MD1-MD3), two exhibited strong and stable interactions between D12 and CS residues (MD1 and MD2). In contrast, during the third simulation (MD3), the substrate began to dissociate from the CS at 164 ns. This run was extended to 600 ns, which permitted us to visualize the translocation of the substrate back and forth between the membrane and the CS, as will be described in the next subsection.

We initially focused on identifying the key interactions governing substrate binding and its coupling to the catalytic site (CS), illustrated in **Fig. 2a-b**. These residues, predominantly hydrophobic in nature, included I267, L263-L264, I133, I131, I54, L305, L308, L342, A339, F163, and I58 (ordered based on the frequency of contacts with the substrate). Additionally, two catalytic and highly conserved histidines (H137 and H167), as well as N134, were observed to persistently coordinate the substrate. The sequence conservation scores computed for these D12-coordinating residues (using Shannon entropy subtracted from the maximum entropy using *ProDy* [27] API; see Methods) showed that H137, H311, H167, N268, Y338, L308 and F163 are highly conserved (see **Supplementary Fig. 1a)**.

**Figure 2.**
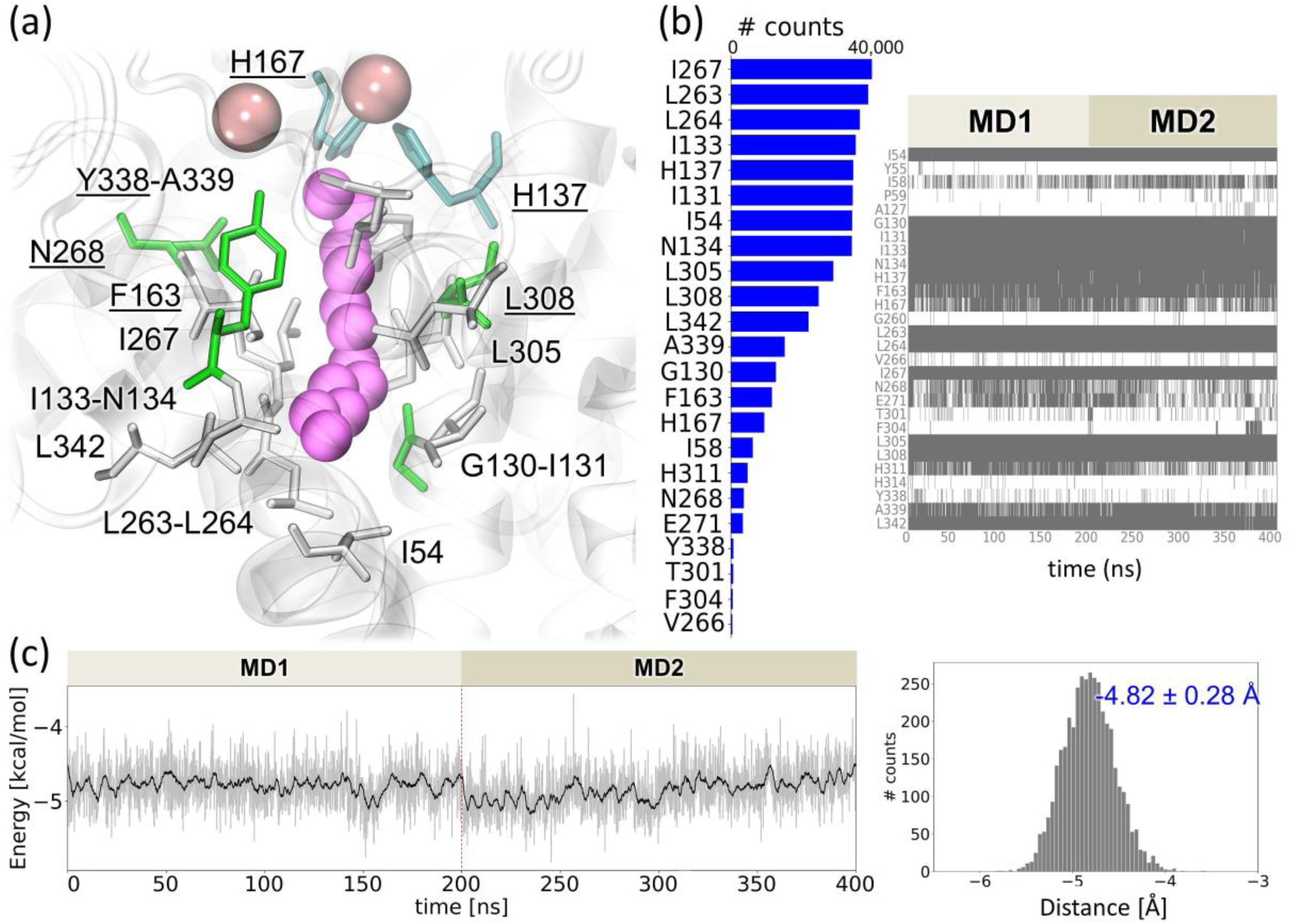
Results from simulations of the AlkB-AlkG complex in the presence of D12, explicit membrane, ions and explicit water. **(a)** A close-up view of the catalytic site and key residues, displayed as *sticks*, that make close contacts with the substrate D12. The underlined amino acids are highly conserved. **(b)** Frequency (*left*) and time evolution (*right*) of atom-atom contacts (within 4 Å). Contact-making residues and the number of contacts they make in two MD runs of 200 ns long each are displayed **(c)** The distribution of D12 binding energy during simulations (*grey*). The black curve shows the moving average using 50 ps windows. The mean binding energy of D12 at the catalytic site is -4.82 ± 0.28 kcal/mol.

The structure and energetics were highly stable during those simulations as can be seen from the time evolution of the binding energy of D12 that remained bound to the CS for 200 ns in runs MD1 and MD2 (**Fig. 2c**). The binding energy fluctuated around -4.8 ± 0.28 kcal/mol (see **Fig. 2c** *histogram*).

### Extended MD simulation unveils a substrate translocation pathway lined by hydrophobic residues that mediate substrate channeling

One of the MD runs, MD3, extended to 600 ns, unveiled a pathway for substrate channeling between the CS and the entrance to the channel located in the membrane as well as the conformational events that facilitate this process (see Supplementary **Movie 1**). **Fig. 3** displays the time evolution of the distance between the substrate and the catalytic iron Fe2 (localized in AlkB and coordinated by histidines and glutamate). The accompanying change in the AlkB - D12 interaction energy is shown in the *bottom*, together with a schematic illustration of the instantaneous positions of the substrate (*pink curly shape*) and Fe ions (three *pink dots*, including one on AlkG). The substrate undergoes a conformational rearrangement between 60 and 90 ns, approximately (**Fig. 3**, *red arrow*) without a significant change in binding energy. This was an early indication of the substrate displacement from the CS that would take place around 190 ns (**Fig. 3**, *black arrow*) to another location and conformation, termed *Intermediate State 1* (*IS1*), where it stayed for almost 150 ns (190-339 ns). *IS1* was approximately 17 Å away from Fe2 and allowed for binding energy comparable to that observed in the CS, leading to a prolonged stay in *IS1*. Then, a subsequent state, *Intermediate State 2* (*IS2*), is visited, bound to a site approximately 4 Å away from *IS1* and ∼1 kcal/mol less favorable (339-478 ns). *IS2* also serves as a gateway to subsequent changes, enabling the access of D12 to the membrane. This state is significantly less stable as the substrate attempts to move toward the membrane. Yet, complete dissociation does not take place. In contrast, a return to *IS1* (with ∼1.5 kcal/mol more favorable energy) occurs for an extended period, following by a dislocation back to CS around 576.5 ns (**Fig. 3**).

**Figure 3.**
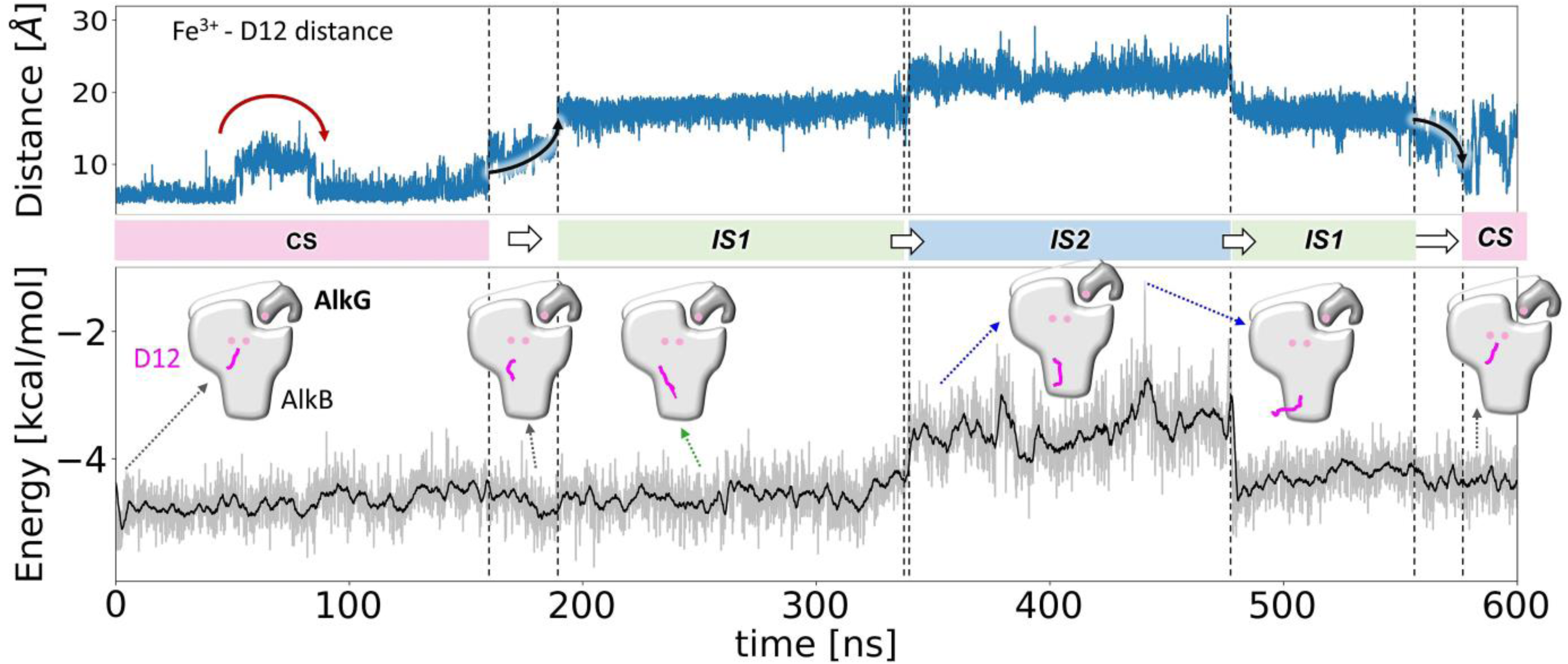
Substrate pathway toward the channel to the catalytic site of FtAlkBG observed in MD3 and Metadynamics simulations. Distance between the closest carbon of D12 and the iron Fe2 (*upper panel*) and corresponding binding energy of D12 binding (*bottom panel*) during MD simulation. *Red-green-blue* bars along the abscissa highlight different stages/locations of D12 during its displacement across the channel; CS – catalytic site, IS1 – intermediate state 1, IS2 – intermediate state 2. Schematic pictures from different stages are included. See also Supplementary **Movie 1.**

During this long trajectory, the substrate nearly exited the protein (at 440.8 ns; **Fig. 3**, *marked by the highest peak*). After that, a notable structural change accompanied by a significant strengthening in binding energy took place around 478 ns, with the substrate translocating back to *IS1* and then back to the CS. Thus, we were able to observe a complete pathway through which the substrate channeled in both directions. Throughout this process, the substrate sampled the same intermediate states (*IS1* and *IS2*) for extended durations, and the binding energy remained at consistent levels.

**Figure 4a** displays the key tunnel-guiding residues, depicted as sticks in *white, green, cyan,* and *orange*. The bar plot in **Fig. 4b** shows their frequency of contacts with D12 where we highlighted in *red* those residues making frequent contacts (during >50% of MD time) at the CS, and in *green* those making less frequent contacts (<50%). Residues guiding the substrate through its transitions between different intermediate states are L53, I27, V128, S124, A127, G50, and P59 in *IS1,* shown by *red sticks* (**Fig. 4a**) and F46, S49, L45, G34, L37, and I33 in *IS2*, shown by *blue sticks* (**Fig. 4a**). The respective groups are marked by *red* and *blue stars* in **Fig. 4b**. Additionally, to more effectively sample the conformational landscape of the channel, we performed metadynamics simulations, which confirmed our MD observations (**Fig. 4b**, *right panel*). The only residues that were not observed in the metadynamics simulations belonged to *IS2,* presumably due to the distance restriction (< 20 Å) imposed between the substrate and Fe2. **Supplementary Fig. 2** displays the detailed time evolution of the interactions observed in both simulations.

**Figure 4.**
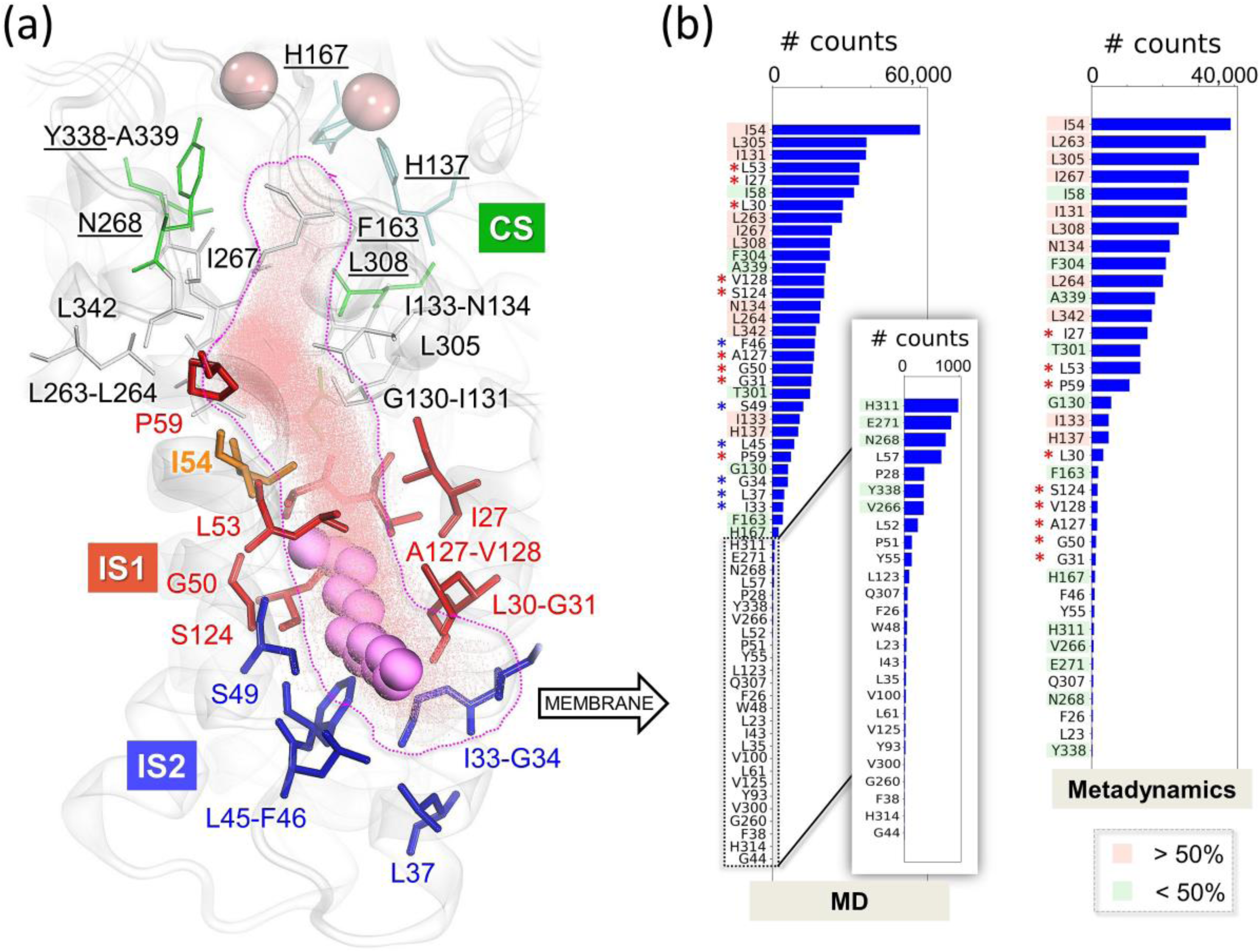
Location of substrate (D12) access channel conducing to the catalytic site of FtAlkBG. **(a)** Close-up view of the catalytic site and substrate translocation channel. Key residues are displayed as *sticks:* those involved in D12 stabilization at the CS are colored *white-green-cyan-orange*; those lining the channel after the departure of D12 from the CS, observed in both MD and metadynamics, are colored *red*; and residues at the end of the channel, at the entrance from the membrane are colored *blue*. The *red* and *blue* residues make frequent contacts with the substrate during the respective intermediates *IS1* and *IS2* (see Fig. 3). The cloud of *pink dots* shows where and how D12 (in *pink space-filling*) traveled during the run MD3 (see **Movie 1**). **(b)** Distribution of contacts (within 4 Å) between FtAlkBG and D12 during 600-ns long MD3 run (*left*) and ∼400-ns long metadynamics simulation (*right*). Residues making the most frequent contacts (>50%) with D12 at the catalytic site are highlighted in *light red*, and those making <50% are highlighted in *light green*. *Red stars* indicate the residues that guide D12 through the channel during *IS1*; *blue stars* indicate those near the entrance (*IS2*).

Further analysis of various conformations sampled by FtAlkBG during MD simulations, revealed the presence of additional channels that were accessible for more than half of the simulation time. As shown in **Fig. 5** (and Supplementary **Movie 2**), a continuous cavity (*blue volume*) was identified, connecting two external entry sites, near R321 and R169 *(orange ovals*) to the CS. These two sites provide access to the CS from the cytosol, and are probably used by O_2_ molecules. At the other end, the channel extends to the membrane-facing entry point near S49 and F46, with a bottleneck regulated by the isomerization of I54 side chain. That exact cavity was used for dislocation from/to the CS by the substrate in MD3 (**Fig. 4** and Supplementary **Movie 1**). Finally, two additional cavities (*green ovals*) were detected; one provided additional volume at the CS (near H171) and the other provided access from the exterior but had no connection to the CS (near W21).

**Figure 5.**
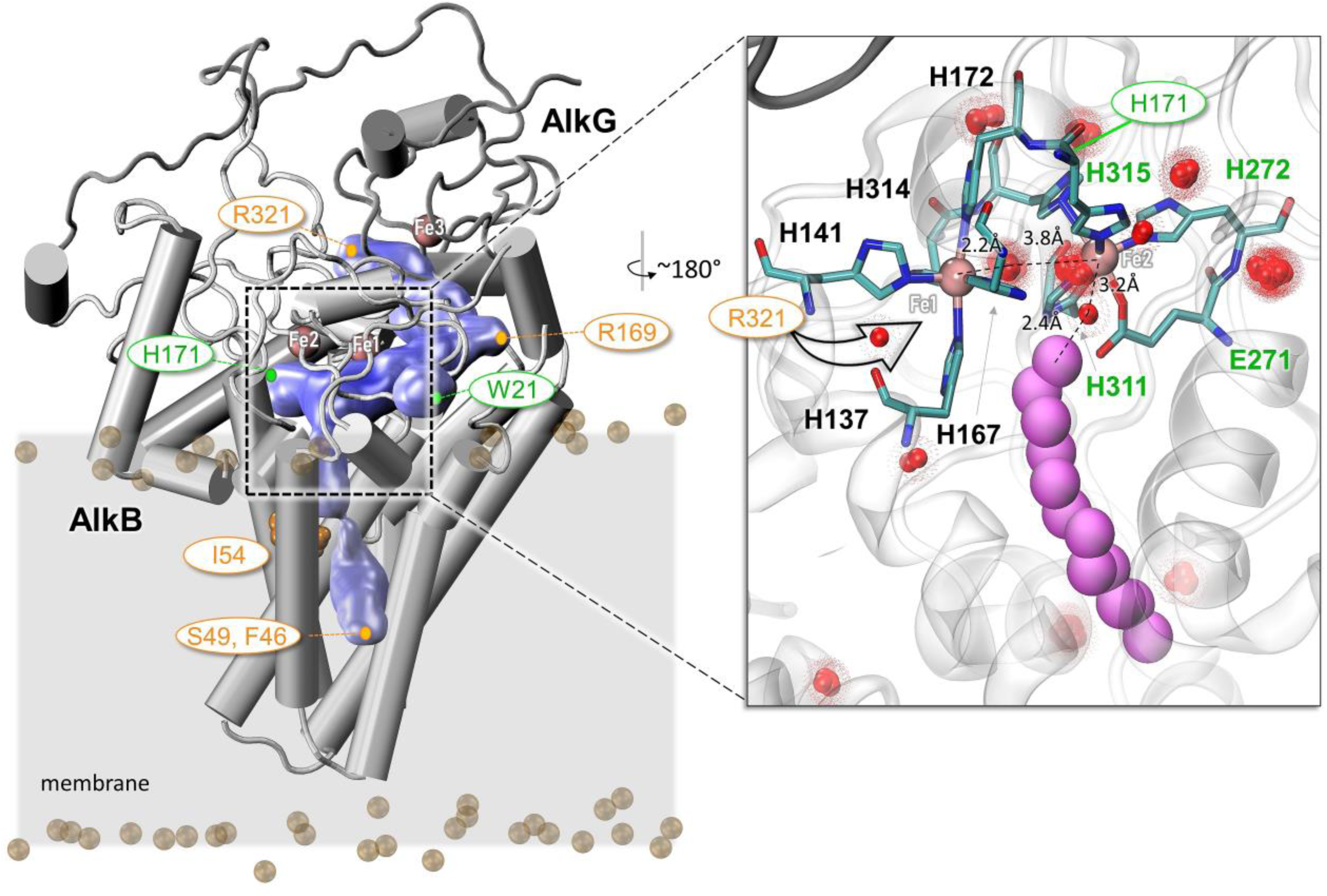
Substrate channeling cavity in FtAlkBG. A major cavity near the catalytic site connecting to substrate translocation channel are shown in *blue surface*, generated by CaviTracer [30] applied to the 600 ns MD3 trajectory. The saturation of *blue* corresponds to the frequency of access. This cavity and channel were accessible (open to the environment) most (> 60%) of the MD simulation. Residues labeled in *orange ovals* denote the entrances from the cytosol (R169 and R321) or the membrane (S49 and F46). *Green ovals* point to partial openings that do not connect to the CS (W21) or to the exterior (H171). *Inset*: Water-mediated associations obtained using WatFinder [29]. Clusters of water are displayed as *red spheres*. The distances between water clusters, irons and/or D12 are shown. The arrow indicates the entrance near R321. See **Movie 2** for a visualization of the cavity from different perspectives.

Knowing how impactful water molecules might be on protein function and/or ligand binding [29], we also examined the location of frequently occurring water bridges (**Fig. 5**, inset). The analysis indicated a substantial presence of water-mediated interactions near the CS. Notably, a high concentration of water molecules was observed between the iron atoms (Fe1–Fe2), and another particularly dense cluster was found near Fe2, along with a few additional water molecules between Fe2 and D12, and at the entrance near R321.

### Alkane binding allosterically consolidates AlkB-AlkG interfacial interactions and the coupling between Fe3 (AlkG) and diiron center (AlkB)

We next examined the stability of the fusion complex FtAlkBG by focusing on the interfacial contacts between AlkB and AlkG (**Fig. 6**). Our analysis unveiled several contacts between highly conserved residues (**Fig. 6a**, *red dots*). Salt bridges in the cytosolic part of the structure persist during 30% of the simulation time (>200 ns, **Fig. 6b**), including D458-R284, D461-R284, D458-K173, D461-R287, R148-D453, D448-R215, R148-E434, and E226-R433 at the interface between AlkB and AlkG (**Fig. 6c**, *red-blue sticks*). Notably, none of these charged residues are conserved (see **Supplementary Fig. 1**).

**Figure 6.**
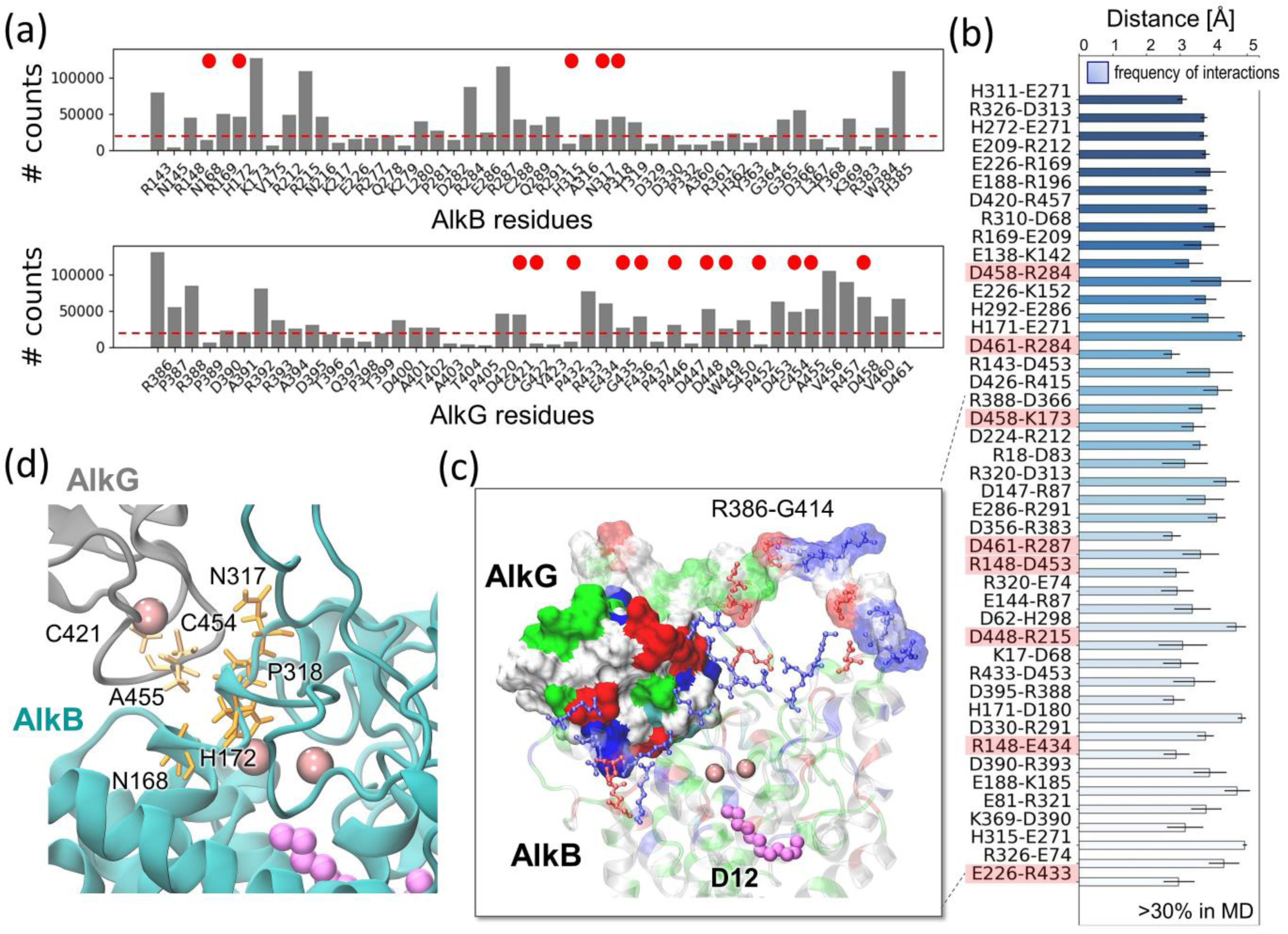
AlkB-AlkG interfacial interactions observed in MD simulations in the presence of D12. **(a)** Interfacial contacts (within 4 Å) between AlkB and AlkG. *Red circles* indicate the highly conserved residues (see **Supplementary Fig. 1**). **(b)** AlkBG residue pairs exhibiting the highest probabilities (>30% total MD time) of forming salt bridges (within 5 Å). Those highlighted in *red* are at the interface between ALkG and AlkB. **(c)** Interfacial salt-bridge forming residues are displayed in *red-blue sticks*. AlkG is in surface representation. The structure is colored by residue type, non-polar (*white*), basic (*blue*), acidic (*red*), and polar (*green*). **(d)** Conserved residues make persistent interfacial contacts at the AlkB-AlkG interface. The analysis is based on all three runs, MD1-MD3.

Overall, considering both the evolutionary conservation of amino acids inferred from Pfam multiple sequence alignments (**Supplementary Fig. 1**) as well as the observed propensities of specific contacts between AlkB and AlkG, the following residues are identified to predominantly underlie the stability of the fusion complex AlkBG: AlkB N168, H172, P318, and N317, and AlkG C421, A455, and C454, illustrated in **Fig. 6d**.

To better understand the impact of substrate binding on AlkBG dynamics, we evaluated the root-mean-square fluctuations (RMSFs) of individual residues in D12-bound and -unbound complexes (**Fig. 7a**). The D12-bound complex showed suppressed fluctuations compared to the unbound at AlkB residues C288-Q289, E179-A182, and N356-A360, and overall AlkG (including salt-bridge forming residues at the interface and the AlkB-AlkG linker R388-G414). These results suggest that substrate binding allosterically consolidated the AlkG-AlkB interfacial interactions as well as the interactions between the linker R388-G414 and neighboring AlkB loop residues C288-Q289 (**Fig. 7a**). In contrast, slightly enhanced fluctuations were observed in the bound form at P73-L80 and A159. Both are away from the D12 binding site, as well as the AlkB-AlkG interface, further supporting the allosteric effect of D12 binding.

**Figure 7.**
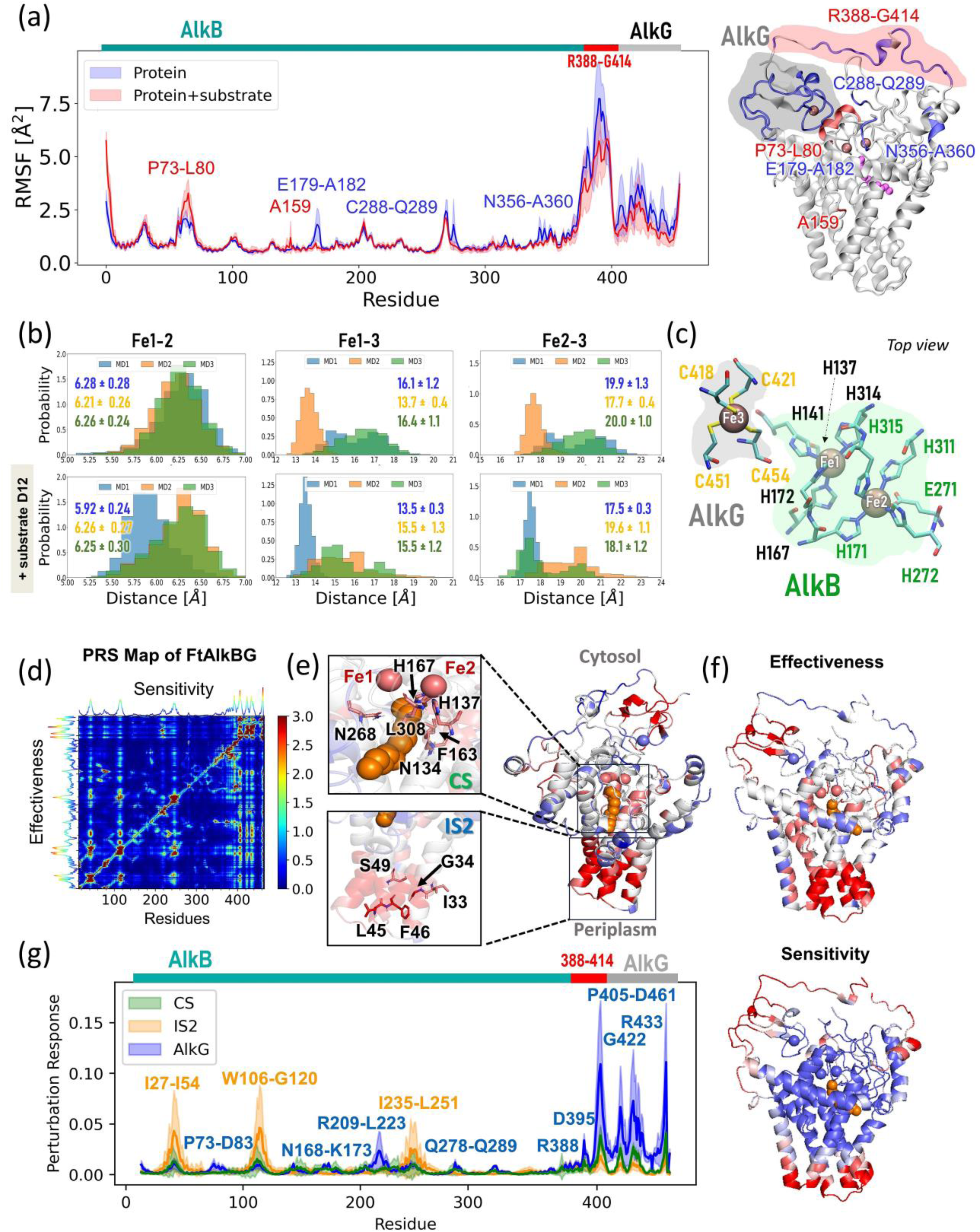
Differences in the dynamics of FtAlkBG with/without the substrate in the catalytic site. **(a)** A comparison of the RMSF profiles of FtAlkBG residues with and without D12. Each curve represents mean values computed from three sets of MD runs with the displayed standard deviation as a shadow (in *red* and *blue*). The cryo-EM structure of FtAlkBG is colored based on the differences in fluctuations between the presence and absence of the D12 substrate. The *grey* and *red* shadows denote the AlkG domain and AlkB-AlkG linker, respectively. *Pink spheres* correspond to irons. **(b)** Histograms showing distances between different iron centers from MD simulations. **(c)** Position and coordination of the three iron-binding sites in FtAlkBG. (**d**) Perturbation Response Scanning (PRS) map of FtAlkBG. **(e)** Effector residues that interact with D12 shown as sticks **(f)** Effectiveness scores for signal transmission mapped onto FtAlkBG structure colored from *red* (highly effective) to *blue* (not effective) **(i)** Comparison of the response curves of residues in the CS, IS2 site, and AlkG in *green*, *orange*, and *blue*, respectively. Each curve represents the mean response of each residue belonging to the particular site, with the standard deviation as a shadow. Peaks of interest are labeled **(h)** AlkBG structure color-coded by sensitivity of residues, from *red* (high) to blue (low).

We further investigated the variation in the distances between the three iron atoms of FtAlkGB, both in the presence and absence of D12 (**Fig. 7b**). The AlkB diiron center is lined by H137, H141, H167, H172, and H314 (coordinating Fe1), and by H171, H272, H311, H315, and E271 (coordinating Fe2). The third iron-binding site, Fe3 or [Fe-4S] [19], located in the AlkG domain, is coordinated by C418, C421, C451, and C454 (**Fig. 7c**) and plays a crucial role in the electron transfer from Fe3 to the diiron center. MD simulations indicate that the Fe1-Fe2 distance remains almost unchanged, around 6.2-6.3 Å across all MD runs with or without D12 (except for one run in the presence of D12 where their average separation is reduced by ∼0.34 Å; see **Fig. 7b**, *left panels*), in line with the 6.1 Å distance observed in the cryo-EM structure [19]. In contrast, the Fe1-Fe3 and Fe2-Fe3 pairs, initially at their respective cryo-EM distances of 13.6 Å and 17.1 Å, experienced broad fluctuations resulting in ∼2Å increases in their average distance (all trajectories; **Fig. 7b** *middle and right panels*). Notably, the strengthening of interfacial interactions induced by D12 binding also brought Fe3 and diiron center closer by ∼0.5 Å on average as can be seen from the comparison of the upper and lower parts of **Fig. 7b** *middle and right panels*.

### Perturbation response scanning analysis elucidates the molecular basis of the allosteric effects induced by substrate binding

We evaluated the overall responses of the enzyme to changes in local interactions using the Perturbation Response Scanning (PRS) method [31, 32]. PRS uses an elastic network model (ENM) representation of the structure. As in information-theoretic approaches, PRS tells us how the application of a force to a given ‘broadcasting’ residue elicits a response (displacement in the equilibrium position) in each of the ‘receiving’ residues[33]. The force may originate from ligand binding, amino acid substitution or other external or internal effects. The procedure scanned over all residue pairs provides a measure of the effectiveness of (perturbed/broadcaster) residues in transmitting signals, and the sensitivity of responding/receiver residues. The response is represented by a color-coded heatmap, the PRS map (**Fig. 7d**), where the perturbed residues are listed along the *y*-axis, and responding residues along the *x*-axis, and the entries in the map are colored from *blue* (low response) to *red* (high response). The curve along the upper abscissa refers to the average response of each residue to perturbation from all others, shortly called the *sensitivity profile* of residues; and that along the left ordinate describes the strength of perturbation exerted by each residue, averaged over all responding ones (*effectiveness profile*). Peaks in the two respective curves are termed sensors and effectors. The ribbon diagrams **Fig. 7e-f** and **7h** are color-coded (*red*, strongest; *blue*, weakest) based on the propensity of residues to serve as effectors and sensors of allosteric signals, respectively.

Examination of the FtAlkBG PRS map reveals several regions with high propensities to serve as effectors. These are composed of sequentially distant but spatially close residues illustrated in **Fig. 7e** insets. One cluster encompasses the residues I33, G34, L45, F46, and S49 making contact with the substrate at early stage of entry from the membrane (*IS2* state; see **Fig. 3**). These residues are therefore distinguished by their enhanced capabilities to communicate perturbations (such as D12 binding) to distal regions, hence the allosteric couplings detected in MD simulations during substrate translocation in the *IS2* state. The second cluster of effectors (L308, N134, F163, N268, together with the iron-coordinating H137 and H167) complements the signal transduction initiated by *IS2* residues.

As to sensor residues, those usually lie in the peripheral regions (**Fig. 7h**). They are manifested by bright (*cyan-to-red*) vertical bands in the PRS map, indicative of their high sensitivity in general. For a clearer assessment of the sensitivity profile of selected regions (CS, *IS2* and AlkG), we generated the sensitivity profiles for those regions, in **Fig. 7g**, using the rows of the map corresponding to these sites. Large peaks tend to correspond to residues with larger RMSF values (**Fig. 7a**), showing that higher flexibility accompanies higher sensitivity. The AlkG response profile shows a small peak at P73-D83, and the CS/AlkG both have another at Q278-Q289, consistent with MD simulations. Perturbations to either the CS or the IS2 sites cause similar responses further demonstrating their strong coupling (**Fig. 7g**). Interestingly, effectors in AlkG also appear to cause a response in *IS2* residues, suggesting that not only does D12 binding stabilize AlkB-AlkG interactions, but that AlkG binding to AlkB may also influence AlkB-D12 interactions. These interactions between the IS2 region, the CS, and AlkG may act as positive feedback that enhances the cooperativity of the structure to drive forward catalysis.

### Cross-correlations between residue movements in the soft modes

To better understand how AlkG and substrate-binding sites are allosterically coupled, we evaluated the cross-correlations between the residues interacting with D12 and those lying at the AlkB-AlkG interface using the Gaussian Network Model (GNM) analysis [34, 35] of soft (energetically favorable) modes of motion. And we further examined the cross-correlations with iron-coordinating residues, AlkG linker, and the residues that exhibited changes in their RMSF upon D12 binding. The results are presented in **Fig. 8**. Panel **a** displays the cross-correlations between residues belonging to different regions, as labeled; the curves along the lower abscissa and left ordinate represent average (absolute) correlations of the particular residues permitting us to identify the residues engaged in strongest cross-correlations (marked by *red dots*), and panel **b** displays the location of the residues distinguished by strong correlations. Cross-correlations range from -1 (fully anticorrelated, coupled but moving in opposite directions) to +1 (fully correlated, moving in the same direction), with 0 corresponding to uncorrelated motions.

**Figure 8.**
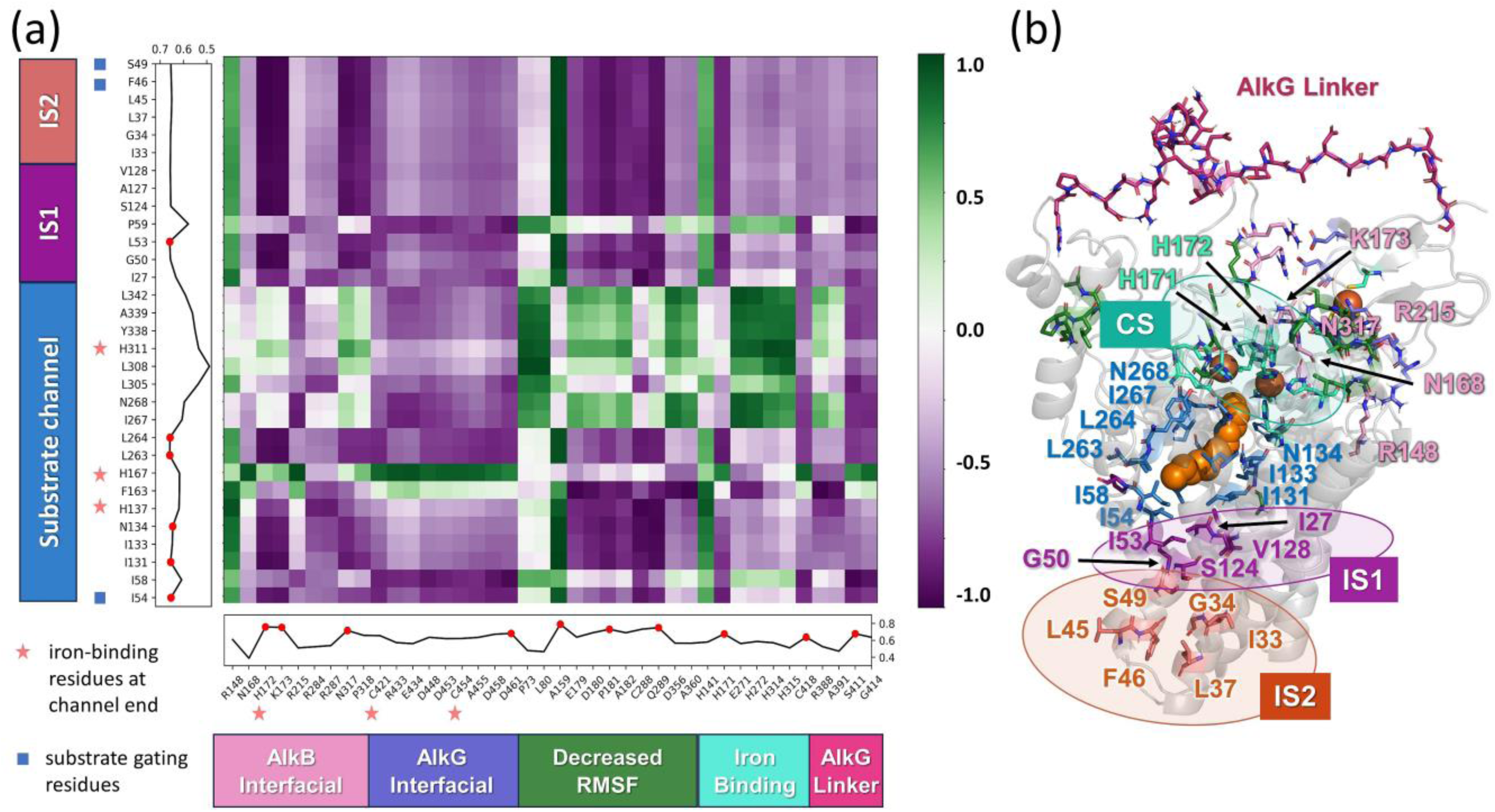
**(a)** Cross-correlation map between different functional sites of FtAlkBG obtained from the first three softest modes predicted by the GNM. Boxes on the x-axis indicate whether residues are part of the AlkB or AlkG interface, and boxes on the y-axis indicate during which state (CS, IS1, or IS2) the residues are interacting with the substrate. Line plots show the RMS cross-correlation values of each residue. **(b)** FtAlkBG structure with residues chosen for cross-correlation analysis displayed as sticks, color-coded the same as the boxes in (**a**), showing the category the residue belongs to.

The cross-correlation map shows confirms that several hydrophobic residues along the channel (e.g. I54, I131, I133, L263, L264; labeled in *blue* in **Fig. 8b**), are strongly correlated with AlkB-AlkG interfacial residues K173, N317 as well as the iron-binding H172, H171 and H315, meaning that these residues are likely responsible for communicating signals from substrate channel to AlkB-AlkG interface through the iron-binding site.While these residues are not in direct contact with the interfacial residues, I131-N134 as well as L263-L264 and A159 serve as a bridge enabling communication; this would also explain the increase in RMSF observed for A159 during MD simulations. As for I54, it is more towards the periplasmic region of FtAlkBG where contacts are made between the transmembrane helices that may be responsible for connecting the dynamics of this distant residue with AlkG (**Fig. 8b**). This also aligns with the PRS data that show that the periplasmic region of FtAlkBG has a cluster of effectors and sensor (**Fig. 7f** and **h**).

Bridging the gap between the D12 binding sites and AlkG lies the diiron site on AlkB (**Fig. 8a-b**). Residues H137, H167, and H311 both interact with D12 and bind iron (**Fig. 8a-b**). The AlkB iron-binding residue H172, as well as the AlkG iron-binding residues C421 and C454 are also part of the AlkB-AlkG interface (**Fig. 8a-b**). This suggests that the iron-binding site plays a critical role in transmitting structural/dynamical changes from D12 binding site to the AlkB-AlkG interface. This idea aligns with results from PRS, which identified H167 and both irons in the diiron complex as effectors (**Fig. 7g**).

A similar phenomenon can be seen for the IS1 and IS2 binding sites. All residues from the IS2 binding site, and most residues in IS1, are strongly anticorrelated with the interfacial and CS/iron-bibnding residues such as H171, H172, K173, N317, and P318, as well as Q289 near the interface. These couplings allow for perturbations caused by D12 binding to be communicated to interfacial residues for enhancing AlkB-AlkG association. Cross-correlation (**Fig. 8a**) and MD simulation data (**Fig. 6**) also indicate that N168, H172, K173, R215, and N317 act as key interfacial residues for the transmission of allosteric signals and for the stabilization of the AlkB-AlkG complex.

### QM/MM models of the active site

We performed hybrid quantum mechanics/molecular mechanics (QM/MM) calculations to investigate the structural features of the non-heme diiron active site of the FtAlkBG and its mechanism of action towards oxygen activation. QM/MM models provide flexibility for the motion of the Fe centers and the coordinating residues and this is reflected in the model starting with no coordinated H_2_O ligands to the Fe centers. Firstly, the Fe1-Fe2 distance decreases to 5.91 Å from 6.2 Å in cryoEM structure for the reduced Fe1(II)-Fe2(II) species. Secondly, geometry optimizations for O_2_ coordination to each of the metal centers lead to formation of a peroxo bridge Fe1(III)-OO-Fe2(III) species instead of individual superoxo Fe^III^OO^•−^ species with an even further reduced Fe1-Fe2 distance of 5.72 Å (**Supplementary Fig. 3).** Formation of Fe1(III)-OO-Fe2(III) species has significant impact on the mechanism for O_2_ and following C-H activation, as it paves the path for formation highly oxidized iron oxo species with O-O bond cleavage [36]. Similar to the Fe1(II)-Fe2(II) case, starting with single water coordination to the active site led to the formation of Fe1(III)-OO-Fe2(III) species as the most stable structure after oxygen coordination, confirming the diferric peroxo species as a critical intermediate. Computational modeling efforts for detailed mechanisms of electron and proton transfer to the FtAlkBG diiron active site and mechanisms of O_2_ and following C-H activation steps are in progress and will be reported elsewhere.

## Conclusion

CryoEM structure resolution of the fusion complex AlkBG enabled the present study to gain a deeper understanding of the time-resolved mechanism of function of this alkane-oxidizing enzyme. Overall, the comparison of the fusion complex in the presence and absence of the alkane (here D12) substrate pointed to the allosteric consolidation of interfacial interactions between AlkB and AlkG upon binding the substrate, also manifested by a rapprochement in the distance between their respective irons Fe1-Fe2 of AlkB and Fe3 of AlkG. Substrate binding and translocation across a long channel, through successive intermediate states *IS2* and *IS1*, is shown to trigger cooperative conformational changes or allosteric effects that facilitate the enzymatic function of the complex. The intrinsic ability of the complex to offer a cavity to accommodate the substrate, as well as a hydrophobic tunnel enabling the channeling of the substrate from a membrane-embedded entry site, is essential to function. Simulations reveal the entry near AlkB F46 and S49, gated by I54, where the hydrophobic tunnel shows a severe constriction. Additional openings of the same cavity to the cytosol suggest paths of entry of O_2_ for the oxygenase activity, while the alkane entry pathway is suggested by simulations as also the exit pathway for the hydroxylated alkane product (the alcohol). The currently identified mechanisms of substrate channeling and allosteric coupling provide new insights into critical sites that could be targeted for producing alternative forms of alkane hydroxylating enzymes and addressing current needs for transforming alkanes to produce energy.

## Methods

### Model preparation and parameterization

In order to determine the full structural model of the alkane 1-monooxygenase AlkBG fusion complex from *Fontimonas thermophila* (FtAlkBG), we used the cryo-EM structure that we have recently resolved (PDB code: 8f6t [19]) and reconstructed the missing R388-G414 linker between AlkB and AlkG using AlphaFold [37] (see **Fig. 1**). Next, we used PPM/OPM server[38] to predict the orientation of the complex in the membrane and CHARMM-GUI[39] to prepare the system with the membrane with the following composition: 50% POPG, 30% POPE, and 20% of cardiolipin molecules (type PMCL2). Further, we parameterized three iron-binding sites of FtAlkGB: two located in AlkB: (i) *Fe2*, coordinated by H171, H272, H311, H315, and E271 (ii) *Fe1*, coordinated by H137, H141, H167, H172, and H314; and one located in AlkG: iron (designated as *Fe3*) coordinated by four cysteines (C418, H421, HC451, C454). Partial charges and geometries for parameterization were obtained from QM/MM calculations. We prepared two systems, with/without dodecane D12. D12 parameters were obtained using SwissParam [40] server and Gaussian (version 16) calculations (DFT B3LYP/6-31(d,p) method).

### Molecular Dynamics simulations

Full-atomic MD simulations were performed using NAMD [22] package and the CHARMM [41] force field, and 2 fs time steps. The system was solvated with explicit water (TIP3P) at physiological salt concentrations. CHARMM force field parameters for the coordination of irons were prepared as described above. For each system, with/without the substrate, we performed 200 ns simulations (three runs per system). One trajectory in the presence of D12 was extended to 600 ns because the substrate moved from the catalytic site and started to approach the channel toward the membrane. The simulated system contained over 205,000 atoms, including those in the lipid bilayer, the water molecules, and KCl ions. The simulation protocol for a membrane was divided into six preparatory simulations before the main run as recommended in CHARMM-GUI protocols [42]. We used our own scripts in ProDy [27] API (including the newly developed *InSty* [43] module) and VMD [44] for analyses and visualization. The binding energies of the substrate were obtained using SMINA [45]. Analysis of water clusters was performed using WatFinder[29] accessible at the ProDy API[27], using the default parameters. Water clusters in MD were identified using the *findClusterCenters()* function with maximum distances between water molecules set to 0.2 Å (dictC) and the minimum number of identified waters set to 10 (numC). The total number of frames was 600 (1 frame per 1 ns of simulation). The cavity/channel predictions were performed using CaviTracer [30] with *calcOverlappingSurfaces()* function.

### Enhanced Sampling Simulations

Metadynamics simulations were conducted using PLUMED plug-in[23] in conjunction with NAMD [22]. In this method, we modified the configurational space of the collective variables (CVs) assigned to the distance between Fe2 and D12 (the closest carbon). The system’s motion along these variables was accelerated by introducing an artificial Gaussian-shaped bias potential to the energy in CV space at each step [46]. This addition aimed to discourage the system from revisiting previously explored states and keeping the simulated system away from the kinetic traps in the potential energy surface. We use Well-Tempered Metadynamics[47] at a temperature of 303.15 K. The data were collected until the system sampled all values of Fe2-D12 distance along the channel. We fixed the maximum value of Fe-D12 distance to 20 Å, and we used the Gaussian height 0.6 and sigma 0.05 with bias factor 12. The total simulation duration was 36 ns.

### QM/MM calculations

The QM/MM model was constructed starting from the coordinates of a representative MD simulation structure. QM/MM optimization was performed using the two-layer ONIOM method with hydrogen link atoms [48], as implemented in Gaussian16 software package. The QM layer was modeled at the B3LYP level of theory [49, 50] using the def2-TZVP basis set [51] for Fe, and the def2-SVP basis set [51] for H, C, N, and O. The QM region was chosen to include the two iron centers and all directly ligated side chains (H137, H141, H167, H171, H172, E271, H272, H311, H314 and H315). Residue E271 was modeled as propanoate and histidines as methylimidazoles in the QM layer [48] using the computed ESP charges. The AMBER force field [52] was used to model the MM region. All the residues within 8 Å of Fe1 and Fe2 centers were allowed to relax during the QM/MM optimization.

### Sequence Analysis using Pfam Data

We analyzed a large set of AlkB (Pfam ID: PF00487) and AlkG (Pfam ID: PF00301) sequences based on a multiple sequence alignment (MSA) retrieved from the Pfam database[53]. The MSA was refined by removing poorly aligned sequences and highly similar sequences that provide redundant data. These filtering criteria (*seqid* = 0.9 and *rowocc* = 0.7) led to a final alignment of 3,649 sequences for AlkG and 24,086 sequences for AlkB. The sequence conservation properties of the filtered MSA were performed using the Evol module [54] of ProDy[27].

### Perturbation Response Scanning and GNM Cross-correlations

The reconstructed FtAlkBG model was used to build an anisotropic network model (ANM) [24]. The ANM was built using the C^α^ atoms and the Fe atoms as nodes. Residues P281-G283, A394-T396, and P466-A467 were reduced using the ReducedModel [55] function from *ProDy* [27]. Perturbation Response Scanning (PRS) was performed using the PRS module [31] of *ProDy* [27]. The GNM analysis of cross-correlations were performed for AlkBG using the cryoEM structure and the *ProDy* interface [27, 56]. GNM nodes were located at the C^α^ atoms and Fe atoms.

## Acknowledgments

This work was supported by the Physical Biosciences Program within the US Department of Energy (DOE), Office of Science, Office of Basic Energy Sciences, Division of Chemical Sciences, Geosciences and Biosciences (grant no KC030402), and utilized computational resources at the Center for Functional Nanomaterials, which is a U.S. Department of Energy Office of Science Facility, and the Scientific Data and Computing Center, a component of the Computational Science Initiative, at Brookhaven National Laboratory under Contract No. DE-SC0012704, by National Institutes of Health grants R01 GM139297 and R01 DK116780 (to I.B), and Polish National Science Centre no. 2022/46/E/ST4/00053 (to K.M.-R.). K.M.-R. would like to thank Dr. Katarzyna Walczewska-Szewc from NCU for her invaluable guidance in performing the metadynamics simulations.

