## Supplemental File for "Substrate binding and channeling allosterically modulate the interactions within the AlkB-AlkG electron transfer complex"

### Supplementary Figures

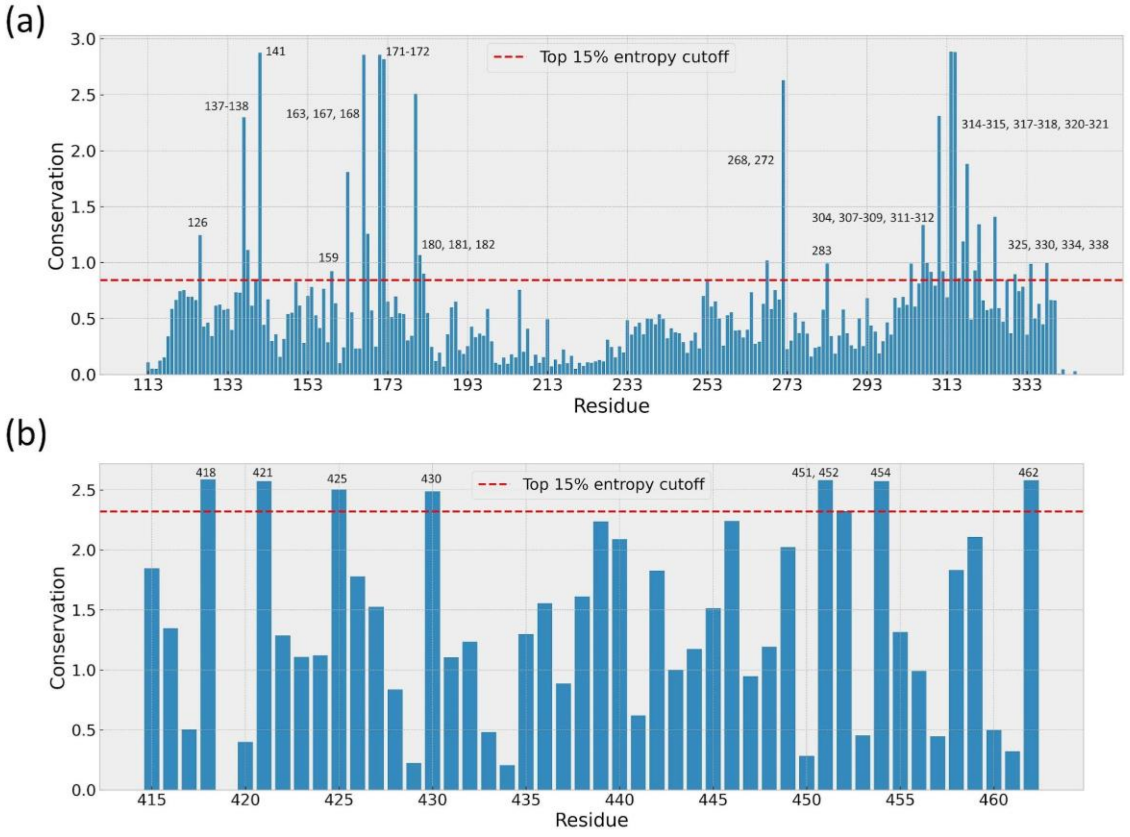


**Supplementary Figure 1. Conservation of AlkB and AlkG proteins.** Conservation propensity of **(a)** AlkB and **(b)** AlkG residues computed based on Shannon entropy subtracted from maximum entropy. The results are based on Pfam multiple sequence alignment data evaluated using the *ProDy* interface. The highest bars correspond to the most conserved residues.

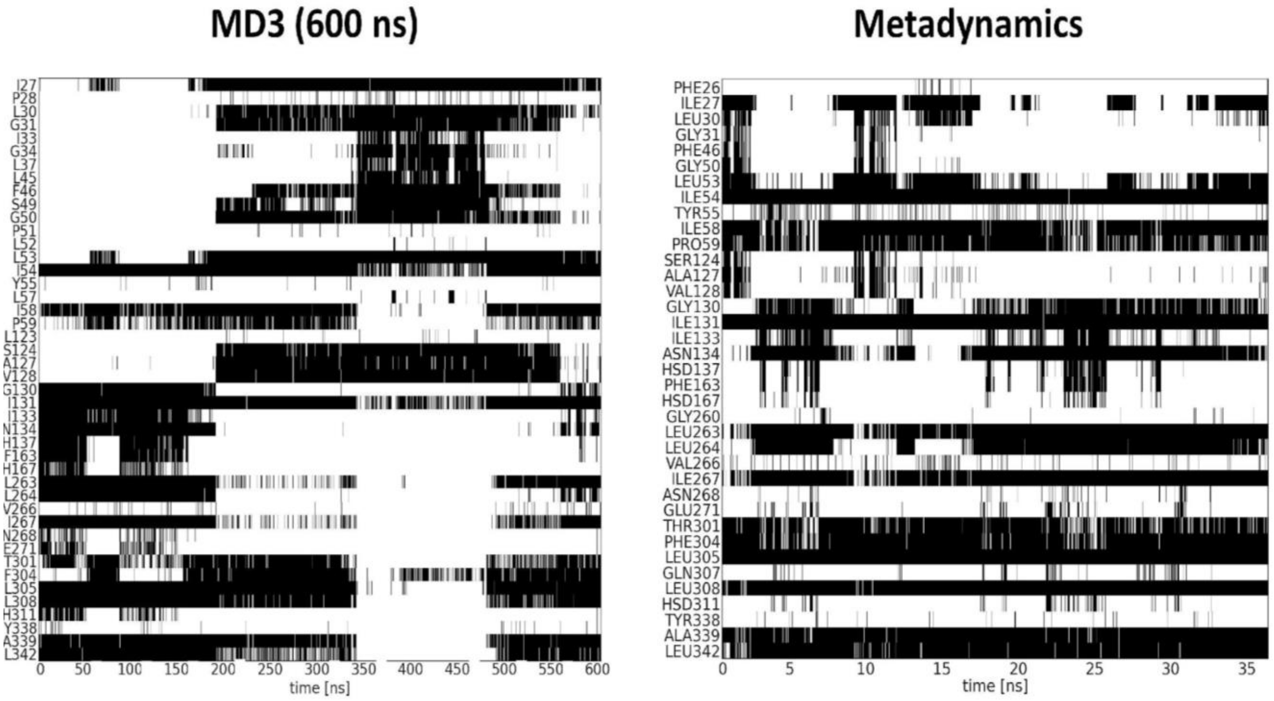


**Supplementary Figure 2. Time evolution of interactions between D12 and AlkB structure observed in classical MD and Metadynamics simulations.** Interactions are indicated by black lines, and persistent interactions appear as black bands

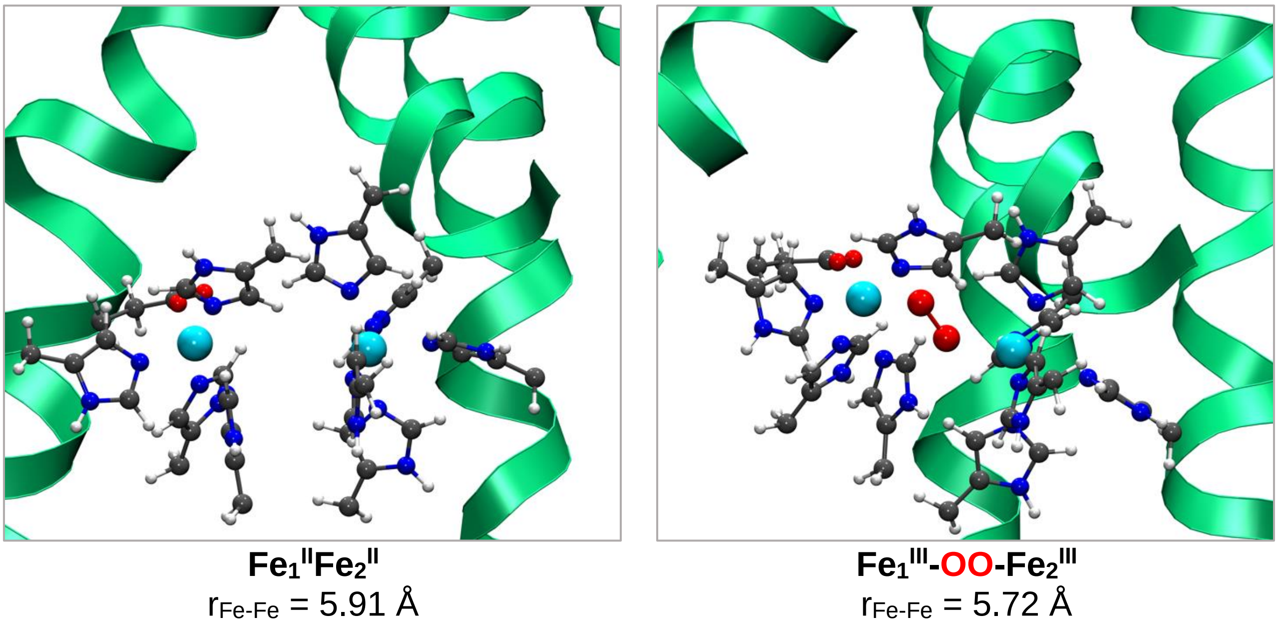


**Supplementary Figure 3. QM/MM optimized structures of active site models of FtAlkBG for $Fe_1^{II}$-$Fe_2^{II}$ (left) and $Fe_1^{III}$-OO-$Fe_2^{III}$ (right).**
